# Plumage colouration differs between offspring raised in natural cavities and nestboxes

**DOI:** 10.1101/2022.08.29.505638

**Authors:** Katarzyna Janas, Irene Di Lecce, Marta Szulkin, Joanna Sudyka

## Abstract

Most of our knowledge on hole-nesting birds, including plumage colouration (an important component of visual signalling), comes from studies on populations breeding in human-provided nestboxes. However, as demonstrated in comparative studies, multiple parameters, such as cavity dimensions and microclimatic conditions, differ between natural and artificial cavities. Despite this, no study so far examined the impact of cavity type on plumage colouration to verify whether extrapolation of results from birds growing in nestboxes is justified. Here, we examined the impact of cavity type – natural cavities vs. nestboxes - on the carotenoid-based colouration of blue tit (*Cyanistes caeruleus*) and great tit (*Parus major*) nestlings. We found clear differences in plumage colouration depending on the type of cavity in which the birds developed. Our study adds to the growing body of evidence confirming that varying properties of natural cavities and nestboxes might influence nestling physiology, leading to phenotypic differences in the long-term.

## Introduction

The progressive anthropogenic impact leads to the disappearance or deterioration of natural habitats (Chamberlain et al., 2009). Besides pollution (Garcia et al., 2017; Querol et al., 2004) and altered microclimatic conditions (Shochat et al., 2006), one of the most serious consequences of this process is the decrease in the availability of suitable breeding sites for wildlife. The shortage of old forest stands is often compensated by providing nestboxes, readily used by some birds, e.g., the great tit (*Parus major*) or the blue tit (*Cyanistes caeruleus*) (Mänd et al., 2005). In a scientific context, nestbox provisioning enables the acquisition of larger sample sizes than in studies of birds breeding in natural cavities (Maziarz et al., 2017; Sudyka et al., 2022a; Wesolowski, 2011). Coupled with facilitated nest monitoring and experimental manipulations (Lambrechts et al., 2010), this leads to a situation where the bulk of ecological research on cavity nesters is conducted on populations breeding in nestboxes. However, natural cavities and nestboxes differ in terms of cavity dimension (Purcell et al., 1997; Sudyka et al., 2022a), microclimatic conditions (Duckworth et al., 2017; Maziarz et al., 2017; Sudyka et al., 2022b), parasite pressure (Fargallo et al., 2001; Wesolowski & Stańska, 2001), and predation risk (Czeszczewik, 2004; Kuitunen & Aleknonis Antanas, 1992; Maziarz et al., 2017; Purcell et al., 1997). Consequently, conclusions stemming from nestbox-breeding populations of birds whose entire evolutionary history is associated with natural cavities, may have limited relevance. This problem has been pointed out in the past (e.g., Møller, 1989, Lambrechts et al. 2010, Wesolowski 2011), which further inspired comparative research on birds breeding in both artificial and natural breeding sites (Czeszczewik, 2004; Norris et al., 2018; Purcell et al., 1997; Sudyka et al., 2022a).

Surprisingly, a topic that has not been examined to date in this context is that of offspring plumage colouration, otherwise remaining in the centre of researchers’ attention (Biard et al., 2017; Drobniak et al., 2013; Isaksson et al., 2006; Peters et al., 2007; Tschirren et al., 2003). To our knowledge, all studies on colouration of hole-nesters come from populations breeding in nestboxes or from birds kept in aviaries, which represents a significant knowledge gap. Despite the fact that the biological meaning of juvenile colouration is still debated, there are indications of its role in parent-offspring communication in both nesting and post-fledging period, informing adults on offspring quality and thus modulating parental provisioning effort (Tanner & Richner, 2008; Galván et al., 2008; Tschirren et al., 2003; Morales & Velando, 2018; García-Campa et al., 2021). Otherwise, it may play a role in establishing social hierarchies between juveniles, act as anti-predatory camouflage in bright deciduous forest (Slagsvold & Lifjeld, 1985; Tschirren et al., 2003), or be the by-product the expression of a genetic correlation with adult colouration (Drobniak et al., 2013).

To date, most work on juvenile colouration comes from nestbox populations of two Paridae species, the blue tit and the great tit, both being important model species in the evolutionary ecology of colour traits. The yellow colouration of blue tit and great tit offspring breast, belly and cheeks is based on carotenoids – lutein and zeaxanthin, deposited in feathers without metabolic modifications (Partali et al., 1987). Since carotenoids cannot be synthesized by birds (Olson & Owens, 1998), their main sources for nestlings are maternal deposition in egg yolk (pre-hatching maternal effects) and pigment-rich food provided by the parents (post-hatching parental effects) (Biard et al., 2005; Isaksson et al., 2006). In adults, carotenoid-based colouration is dependent on their foraging abilities and might thus be a good predictor of their parental quality (García-Navas et al., 2012). Accordingly, blue tit females breast colouration was shown to be positively related with clutch size and fledging success (Doutrelant et al., 2008). Moreover, both breast and UV-blue crown colouration are important in resource allocation, as they are used by a parent to adjust offspring provisioning rates according to the perceived quality of their partner (Limbourg et al., 2013; Mahr et al., 2012).

In this comparative study, we examined the effect of cavity type – natural cavities vs nestboxes – on offspring plumage colouration, and specifically - carotenoid chroma and brightness of blue tit and great tit nestlings breast feathers. Carotenoid chroma and brightness, as determined by distinct properties of the feather, are thought to convey different information about the signal emitter (Shawkey & Hill, 2005). Carotenoid chroma reflects the amount of pigment deposited in the feather (Andersson & Prager, 2006), while brightness is supposed to indicate the quality of reflective keratin structure and might depend on the degree of feather abrasion and the trace amounts of melanin (Isaksson et al., 2008; Shawkey & Hill, 2005).

Here, we used data from two breeding seasons collected in an ecologically homogeneous old-growth forest, where one plot was supplied with nestboxes, while the other, closely situated, offered only natural cavities as breeding sites. This created a unique, quasi-experimental setup that allowed to objectively infer the consequences of breeding in human-provided nests. To ensure that nestling colouration was not biased by preference of differentially-coloured parents towards a particular cavity type, we also examined adult’s colouration in relation to the type of cavity they bred in.

## Methods

### Study site and sampling

The study was conducted in two consecutive field seasons (2018-2019) in Bielany Forest (ca. 150 ha), a remnant of the European primeval forest. Data were collected within two study plots, further referred to as “natural cavity” and “nestbox” plot, with edges separated by min. 200 meters. Both plots are covered by an oak-hornbeam forest (*Tilio-Carpinetum*), with some of the oak stands reaching 300-400 years (Pawlat-Zawrzykraj et al. 2021). Importantly, the small distance ascertained that the plots were functionally identical as they shared the same environmental patch. Accordingly, Sudyka et al. (2022a) found no difference in food availability between study plots in both field seasons, assesses by widely used method of frass fall collection (Wesolowski & Rowiński, 2014). Ambient temperature and humidity as well as anthropogenic factors (noise and air pollution, measured as PM 2.5 concentration) were found to be uniform between study plots (Sudyka et al., 2022b). Furthermore, the risk of predation does not vary at this scale, for example in case of martens such a distance is a territory of one individual/pair (Zalewski & Jędrzejewski 2006).

Nest searches in the natural cavity plot and nestbox monitoring started at the beginning of April. In nestlings, we sampled breast feathers ca. 14 days after hatching (hatching = day 0) and performed basic biometric measurements. In adult blue tits, we sampled crown and breast feathers, while from great tits, only breast feathers. For more details on study site and sampling, sample size (Table S1) and for further details on the contrasting biology of these two species when breeding in natural cavities and nestboxes, please refer to ESM and to (Sudyka et al., 2022a, 2022b).

### Plumage reflectance measurements

We measured reflectance using a spectrophotometer USB400 (Ocean Optics) with 300–700 nm range, a xenon pulsed light source. Before the onset of measurements and every 20 minutes, we took a reference scan of the white standard (Spectralon®). From each sample we took 10 reads with the probe held perpendicular to the feather surface [as in (Janas et al., 2018)]. Raw spectra were averaged and processed in the *pavo* package (Maia et al., 2019) in R. We calculated: total brightness quantified as a sum of reflectance values over all wavelengths (hereafter brightness), and two measures of chroma: carotenoid chroma and UV chroma (Montgomerie, 2009). Carotenoid chroma: 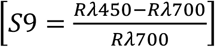 is the relative reflectance in the region of the spectrum with maximum carotenoid absorbance (R_λ450_), thus it constitutes the best approximation of carotenoid content in the feathers. UV chroma was calculated as a total reflectance in between 320-400 nm divided by total reflectance (brightness). Carotenoid chroma and UV chroma of feathers with carotenoid-based colouration, i.e., breast feathers, are strongly, negatively correlated, and both are independent of brightness (Figure S2).

### Statistical analysis

We checked the distribution of colour variables by visual inspection of diagnostic plots. All of them were normally distributed, thus to detect outliers we applied generalized extreme Studentized deviate test (ESD), with type I error = 0.05 (Rosner, 1983). All records identified by the Rosner’s test as outliers were removed from the main analysis, so the exact sample sizes differ among the models. All colour variables (together with clutch size and lay date) were scaled to zero mean and unit standard deviation for clarity of parameter estimates.

To test for differences in colour parameters of blue tit and great tit nestlings raised in natural cavities and nestboxes, we fitted general linear mixed models, separately for brightness and carotenoid chroma as response. To account for the impact of common rearing environment, we fitted a unique nest ID (nest ID with a year code, hereafter: nest ID) as a random term. Full models included cavity type (natural cavities or nestboxes), parental colour metrics and parental age (categorized as 2nd year of life and older, (Svensson, 1994), laying date, clutch size, nestlings sex, nestlings mass and year of study, as well as their interaction with cavity type (Table S2 and Table S3 including outliers). In the primary version of full models, we also tested for the quadratic term of laying date (as food availability is distributed non-linearly across a breeding season (Wesołowski & Rowiński, 2014). However, as nonsignificant, it was dropped from all models. In the final models, the following variables of biological relevance were always retained: cavity type, parental colour metrics (to quantify parent-offspring correlations), nestling sex and year (differences in weather conditions and food availability between two study seasons, (Sudyka et al., 2022a). Other factors (parental age, lay date, clutch size, nestling mass), if nonsignificant, were removed from the models in backward elimination procedure (Quinn & Keough, 2022). We controlled for multicollinearity of predictors in final models by checking the variance inflation factor (VIF) following (Zuur et al., 2010), which was lower than 2 in all cases. All statistical analysis were performed in R (version 4.1.1, (R Core Team, 2021)).

## Results

There were clear differences in nestling plumage colour composition depending on whether they developed in nestboxes or in natural cavities. Carotenoid chroma was higher in blue tit nestlings reared in nestboxes relative to those reared in natural cavities (Estimate = 0.497, p-value = 0.013, n = 452, Figure 1b), and it was also found to be negatively related to clutch size (Table 1, Figure 2a). Both brightness and carotenoid chroma differed depending on blue tit nestling sex, as males displayed lower brightness and higher carotenoid chroma (Table 1). There was also a tendency for blue tits raised in nestboxes to have higher breast brightness than those from natural cavities (Estimate = 0.268, p-value = 0.059, n = 450, Figure 1a).

**Table 1.**
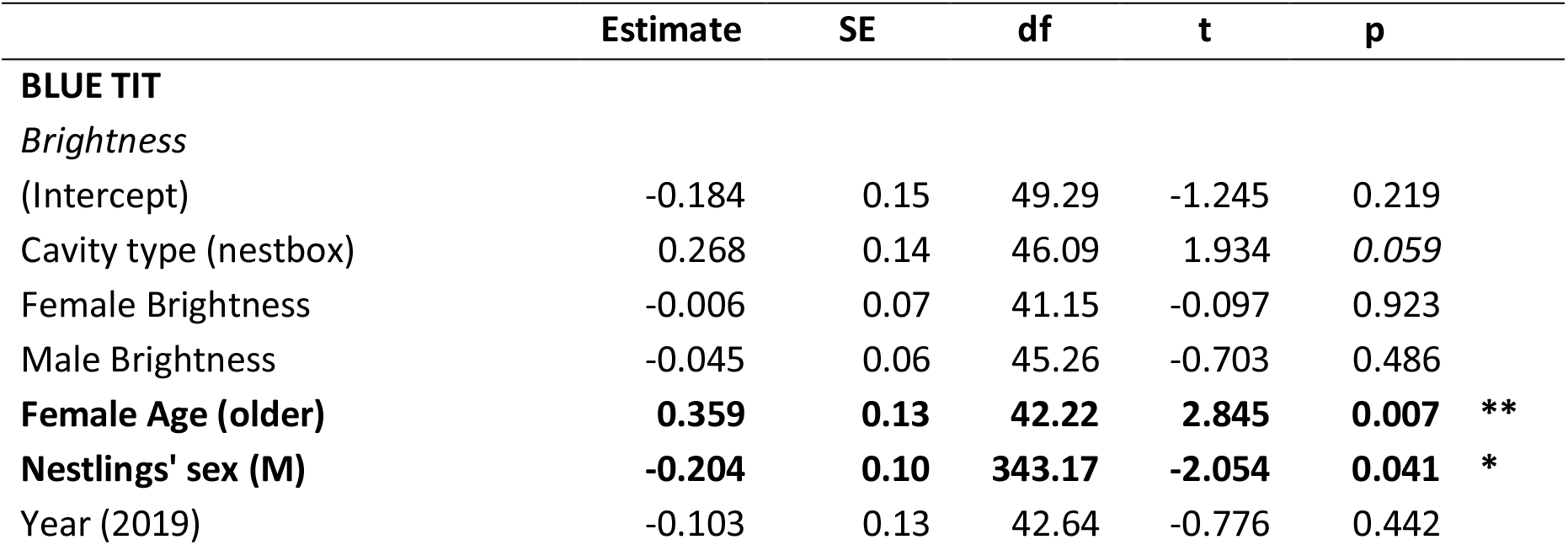
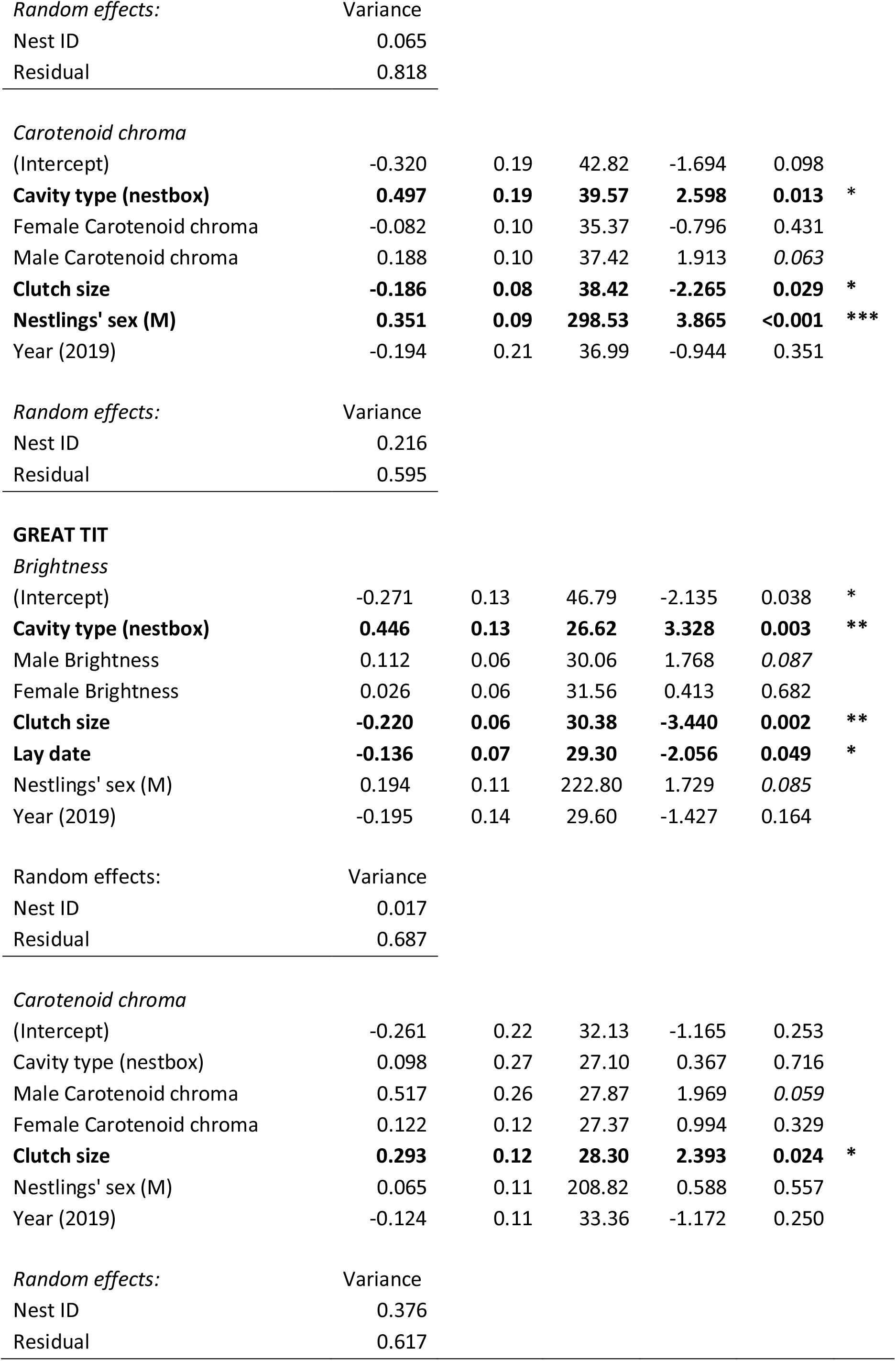
Final models testing the effects of cavity type (natural cavity set as reference value), parental plumage characteristics, age (first year breeding bird or older), nestling sex, and year on plumage traits of blue tit and great tit nestlings. All colour variables, clutch size and lay date were scaled to zero mean and unit SD. P<0.1 are given in italics, *P<0.05, **P<0.01, ***P<0.001.

|  | Estimate | SE | df | t | p |
| --- | --- | --- | --- | --- | --- |
| **BLUE TIT** |  |  |  |  |  |
| *Brightness* |  |  |  |  |  |
| (Intercept) | -0.184 | 0.15 | 49.29 | -1.245 | 0.219 |
| Cavity type (nestbox) | 0.268 | 0.14 | 46.09 | 1.934 | *0.059* |
| Female Brightness | -0.006 | 0.07 | 41.15 | -0.097 | 0.923 |
| Male Brightness | -0.045 | 0.06 | 45.26 | -0.703 | 0.486 |
| **Female Age (older)** | **0.359** | **0.13** | **42.22** | **2.845** | **0.007** ** |
| **Nestlings' sex (M)** | **-0.204** | **0.10** | **343.17** | **-2.054** | **0.041** * |
| Year (2019) | -0.103 | 0.13 | 42.64 | -0.776 | 0.442 |

|  |  |
| --- | --- |
| *Random effects:* | Variance |
| Nest ID | 0.065 |
| Residual | 0.818 |

*Carotenoid chroma*
|  |  |  |  |  |  |  |
| --- | --- | --- | --- | --- | --- | --- |
| (Intercept) | -0.320 | 0.19 | 42.82 | -1.694 | 0.098 |  |
| **Cavity type (nestbox)** | **0.497** | **0.19** | **39.57** | **2.598** | **0.013** | * |
| Female Carotenoid chroma | -0.082 | 0.10 | 35.37 | -0.796 | 0.431 |  |
| Male Carotenoid chroma | 0.188 | 0.10 | 37.42 | 1.913 | 0.063 |  |
| **Clutch size** | **-0.186** | **0.08** | **38.42** | **-2.265** | **0.029** | * |
| **Nestlings' sex (M)** | **0.351** | **0.09** | **298.53** | **3.865** | **<0.001** | *** |
| Year (2019) | -0.194 | 0.21 | 36.99 | -0.944 | 0.351 |  |

|  |  |
| --- | --- |
| *Random effects:* | Variance |
| Nest ID | 0.216 |
| Residual | 0.595 |

*Brightness*
|  |  |  |  |  |  |  |
| --- | --- | --- | --- | --- | --- | --- |
| (Intercept) | -0.271 | 0.13 | 46.79 | -2.135 | 0.038 | * |
| **Cavity type (nestbox)** | **0.446** | **0.13** | **26.62** | **3.328** | **0.003** | ** |
| Male Brightness | 0.112 | 0.06 | 30.06 | 1.768 | 0.087 |  |
| Female Brightness | 0.026 | 0.06 | 31.56 | 0.413 | 0.682 |  |
| **Clutch size** | **-0.220** | **0.06** | **30.38** | **-3.440** | **0.002** | ** |
| **Lay date** | **-0.136** | **0.07** | **29.30** | **-2.056** | **0.049** | * |
| Nestlings' sex (M) | 0.194 | 0.11 | 222.80 | 1.729 | 0.085 |  |
| Year (2019) | -0.195 | 0.14 | 29.60 | -1.427 | 0.164 |  |

|  |  |
| --- | --- |
| *Random effects:* | Variance |
| Nest ID | 0.017 |
| Residual | 0.687 |

*Carotenoid chroma*
|  |  |  |  |  |  |  |
| --- | --- | --- | --- | --- | --- | --- |
| (Intercept) | -0.261 | 0.22 | 32.13 | -1.165 | 0.253 |  |
| Cavity type (nestbox) | 0.098 | 0.27 | 27.10 | 0.367 | 0.716 |  |
| Male Carotenoid chroma | 0.517 | 0.26 | 27.87 | 1.969 | 0.059 |  |
| Female Carotenoid chroma | 0.122 | 0.12 | 27.37 | 0.994 | 0.329 |  |
| **Clutch size** | **0.293** | **0.12** | **28.30** | **2.393** | **0.024** | * |
| Nestlings' sex (M) | 0.065 | 0.11 | 208.82 | 0.588 | 0.557 |  |
| Year (2019) | -0.124 | 0.11 | 33.36 | -1.172 | 0.250 |  |

|  |  |
| --- | --- |
| *Random effects:* | Variance |
| Nest ID | 0.376 |
| Residual | 0.617 |

**Figure 1.**
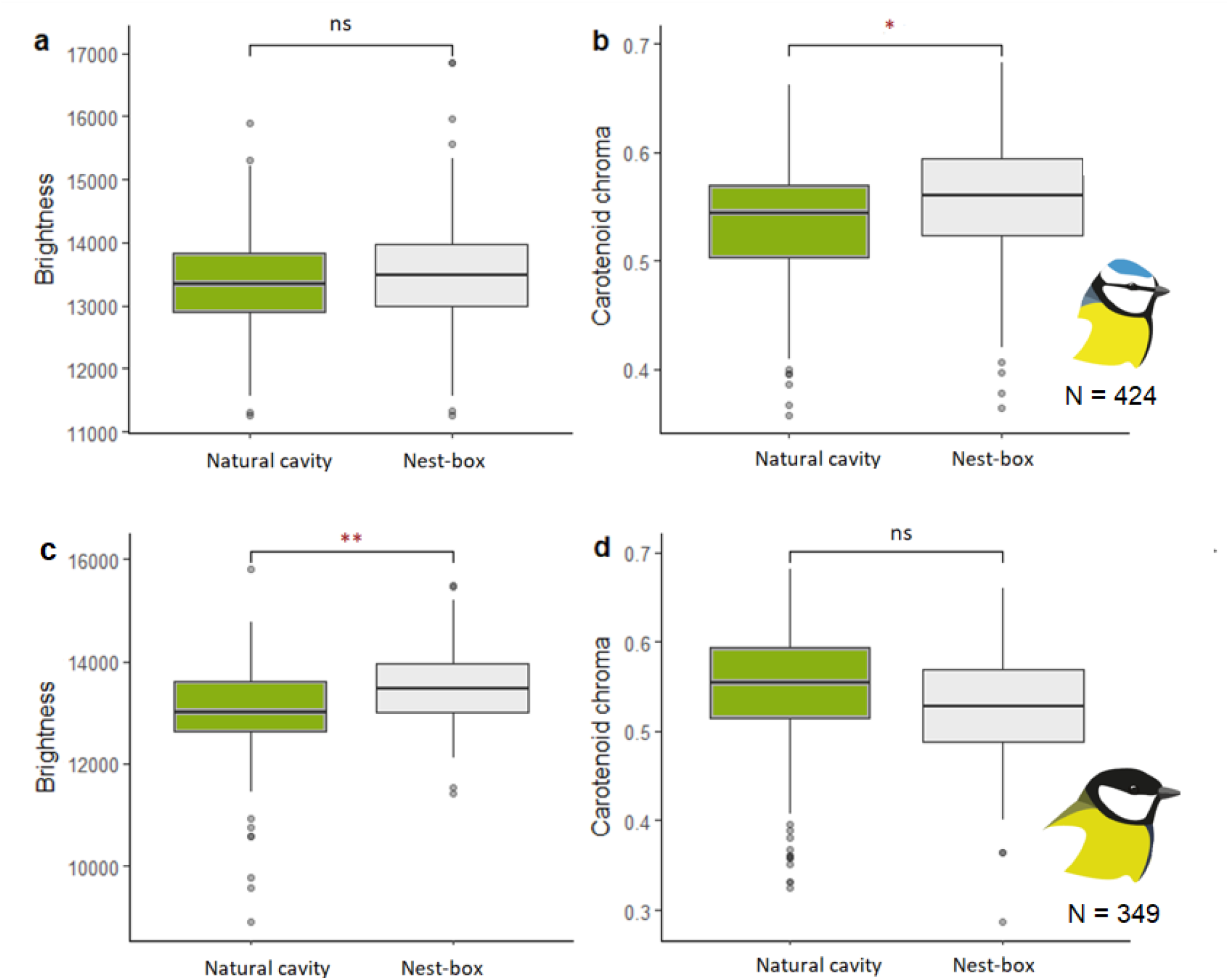
Differences in colouration of blue tit (a,b) and great tit (c,d) nestlings. Horizontal bars denote raw data median, whiskers indicate minimum and maximum values, green and light grey colours denote, respectively, nestlings raised in natural nests and nestboxes. Asterisks indicate significance levels for ‘cavity type’ factor in models presented in Table 1. *P<0.05, **P<0.01.

**Figure 2.**
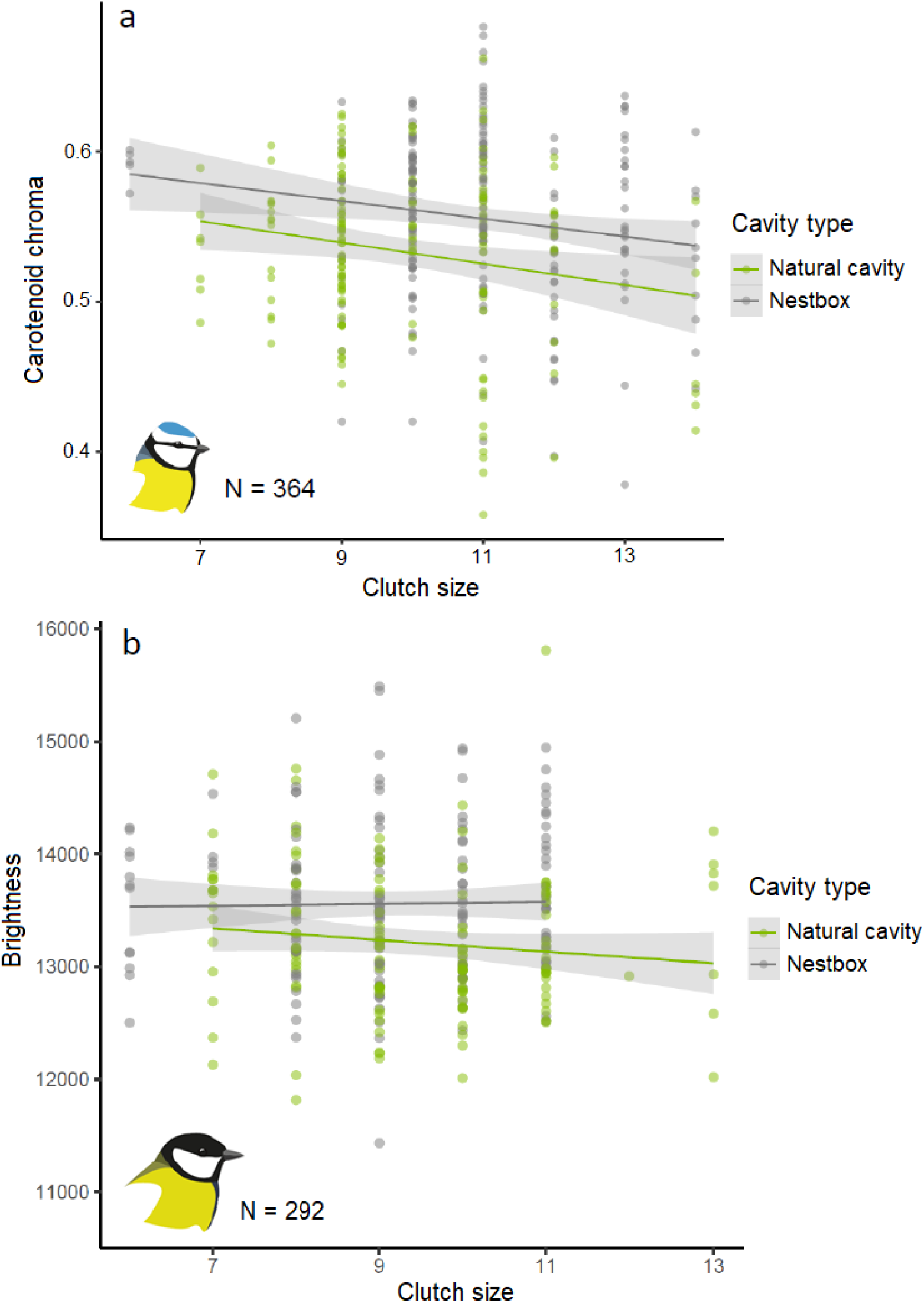
Relationship between carotenoid chroma of blue tit (a) and great tit (b) nestlings and clutch size (Table 1). Green points and line represent nestlings from natural cavities and grey points and line denote nestlings raised in nestboxes. Raw data with outliers removed.

In great tits breast brightness was higher in nestlings raised in nestboxes relative to natural cavities (Estimate = 0.446, p-value = 0.003, n = 359, Figure 1c). It was also found to be negatively associated with both laying date and clutch size (Table 1). In contrast to blue tits, the expression of carotenoid chroma did not differ between great tit nestlings raised in natural cavities and nestboxes (Table 1, Figure 1d). Additionally, great tit nestlings were heavier 14 days after hatching when raised in nestboxes than in natural cavities (t = -5.796, df = 394.71, p< 0.001), but blue tit nestlings mass was uniform between cavity types (t = 0.858, df = 470.98, p-value = 0.391).

Interestingly, in both species we observed a borderline significant (0.05<p<0.1), positive relationship between nestling and father carotenoid chroma (Table 1).

As expected, there were no differences in adult blue tit crown brightness, breast brightness and carotenoid chroma when breeding in natural cavities and nestboxes (Table S4, Figure S3). Similarly, adult great tit breast brightness did not differ depending on cavity type, but there was a significant interaction between cavity type and year for carotenoid chroma (Table S4, Figure S4).

## Discussion

This study reveals differences in juvenile plumage colouration depending on whether birds were raised in natural cavities or nestboxes. By comparing plumage colouration of juveniles raised in different cavity types, we found carotenoid chroma to be significantly higher in nestbox-reared blue tits, and brightness to be higher in nestbox-reared great tits (with a similar tendency for blue tits) when compared to natural cavities.

The juvenile, cavity-type specific differences in carotenoid chroma in blue tits but not in great tits may stem from the disparities in reproductive ecology and sensitivity to conditions in the breeding cavity (Corsini et al., 2017; Sudyka et al., 2022a; van Balen, 2002; van Balen & Potting, 1990). Sudyka et al. (2022a) reported that in the same forest, blue tits breeding in nestboxes had lower hatching and fledging success, spent more time in the nest, and fledged later than those from natural cavities. The negative relationship between clutch size and carotenoid chroma observed in this study are congruent with evidence from previous experimental studies of brood size reduction (Jacot et al., 2010; Tschirren et al., 2003), suggesting that while caring for smaller broods, blue tit parents can provide more carotenoid-rich food to their nestlings. In contrast, in great tits, neither carotenoid chroma of nestlings nor parameters of reproductive success (Sudyka et al. 2022a) differed between birds from natural cavities and nestboxes. However, great tit nestlings raised in nestboxes expressed higher feather brightness compared to the ones reared in natural cavities. This could indicate a better quality of feather structure when reared in nestboxes, which is also in agreement with higher great tit body mass 14 days after hatching relative to natural cavities (Figure S5).

Since the effect sizes of differences observed in our study were relatively small, we are careful in interpreting our results in the context of consequences for parents-offspring signalling. However, in line with previous research, we found the presence of sexual dichromatism (both chroma and brightness) in blue tit nestling plumage (e.g., Jacot & Kempenaers, 2007) and lack thereof in great tits (Isaksson et al., 2008), suggesting a possible difference in the signalling role of this trait between species. Although still under debate, growing evidence points to a signalling function of nestling colouration in the blue tit (García-Campa et al., 2021; Morales & Velando, 2018), while in the great tit there is no evidence of parental preference towards more coloured nestlings (Tschirren et al., 2005).

In contrast to limited information on the signalling function of juvenile colouration, the role of adult plumage colouration is relatively better understood. Previous studies reported that feather colouration, especially in carotenoid-based traits, might signal foraging abilities and parental effort (Germain et al., 2010; Senar et al., 2002). Accordingly, it was showed that breast carotenoid chroma in blue tits of both sexes was positively related with provisioning rates (García-Navas et al., 2012), while great tit nestling colouration was correlated with the rearing father’s chroma (Fitze et al., 2003; Isaksson et al., 2006). In line with this, we found in both species a trend at the margin of statistical significance for a positive relationship between nestling and father carotenoid chroma (Table 1). This may be because in the first days after hatching (critical for accumulating the optimal amount of carotenoids), males are the main food providers, while females spend most time in the nest warming the nestlings. Moreover, we found no evidence for preference of high-quality adults (in terms of plumage colouration) towards a specific type of cavity in either species (Table S4), and consequently assume that differences in nestling plumage reported here are not driven by differences in plumage-based parental quality.

Finally, a potential limitation of our study might be the spatial separation of the study plots with natural cavities and nestboxes. A previous study (Lõhmus & Remm, 2005) suggested differential competition as well as quality-dependent and species-specific preference towards cavity type in our model species if both types are offered on the same site. Natural cavities are not randomly or evenly distributed within a study site like nestboxes are, thus it is not possible to sample them at random locations within a site to overcome this difficulty. Thus, following previous studies (Czeszczewik, 2004; Mitrus, 2003; Norris et al., 2018) and to ensure random nest site choice we decided to keep the plots slightly separated. It is unlikely that habitat varies in such a small spatial scale, and we confirmed this lack of differences between study plots (see Methods), most importantly in the availability of caterpillars, a crucial factor for determining the intensity of carotenoid-based plumage colouration (Sudyka et al. 2022a). To the best of our knowledge, the potential uncertainty stemming from the slight spatial separation of the plots outweighs the much riskier interspersing of the cavity types and resulting parental-quality bias.

To conclude, our results add to the growing body of evidence highlighting the fact that differences between natural cavities and nestboxes can influence nestling physiology and development. We acknowledge that due to the difficulty of sampling in natural cavities, the research conducted on birds breeding in nestboxes will remain the main source of knowledge on the reproductive biology of secondary cavity nesters. Yet, given the clear differences in nestling colouration reported in offspring reared in nestboxes as opposed to natural cavities, we advocate for further research, particularly focused on the long-term biological consequences of these differences, in terms of survival and future reproductive success.

## Acknowledgements

We thank Patryk Rowiński and numerous assistants for help in fieldwork. We are grateful to Adrian Surmacki and Szymon Drobniak for providing the spectrophotometer and technical support. The study was supported by NCN grant no. OPUS 2016/21/B/NZ8/03082 awarded to M.S.

## Ethics statement

The research was conducted under the permits of RDOŚ, for the natural cavity plot: WPN-I.6401.80.2017. LM, WPN-I.6205.53.2017.AS, and for the nest-box plot: WPN-I.6401.515.2017.KZ and WPN-I.6205.227.2017.AS, as well as Lasy Miejskie – Warszawa – permit no. LM-W.LO.400.88.2017. DC1460.

## Statement of Authorship

K.J.: conceptualization, investigation, data curation, formal analysis, writing—original draft and writing— review and editing; I.D.L.: sample collection, investigation, writing—review and editing; M.S.; funding acquisition, supervision, project administration, writing—review and editing J.S.: conceptualization, sample collection, data curation, formal analysis, investigation, funding acquisition, supervision, writing—review and editing. All authors approved the final version of the manuscript and agreed to be held accountable for the work performed therein.

