## Supplementary material for "Plumage colouration differs between offspring raised in natural cavities and nestboxes"

**Study species**

Blue tits (*Cyanistes caeruleus,* Linnaeus, 1758) and great tits (*Parus major,* Linnaeus, 1758) are small, secondary-cavity nesters, commonly occurring throughout Europe. They preferentially breed in natural cavities with slit-shaped openings in living trees, (formed as a result of wood decay rather than woodpeckers activity) especially in Hornbeam *Carpinus betulus*, Alder *Alnus glutinosa* and Small-leaved lime *Tilia cordata* (Maziarz et al., 2015; Wesołowski & Rowiński, 2012). However, they readily breed in nestboxes, which makes them a great model species in evolutionary ecology (Stenning 2018). Both blue tits and great tits are socially monogamous, with bi-parental care and a relatively high rate of extra-pair copulations (Charmantier et al. 2004). Despite a smaller body size (on average ~11 g, compared to ~17 g in great tit), blue tits lay bigger clutches, with an average number of 12 eggs, compared to 9 in great tits, and they usually start incubation earlier. In both species, incubation lasts around two weeks, but the length of this period might be modified in urban habitats or as a result of extreme weather conditions (Corsini et al., 2017; Glądalski et al., 2018). Nestlings are fed mainly with caterpillars (e.g. the winter moth *Operophthera brumata* L.), which are also the main source of carotenoids: lutein and zeaxanthin (Perrins, 1991; Isaksson et al., 2008). Fledging usually takes place 17-22 days after hatching and most fledglings leave the nest within one day (Stenning 2018). From mid-July to October, juveniles undergo partial postjuvenile moult, which in blue tits includes all body feathers, greater, median and lesser coverts, and might also include alula, carpal covert, tertials and two central tail feathers (Demongin 2016). The pattern of postjuvenile moult in great tits is similar, yet it can include some inner secondaries and some to all tail feathers. Adults of both species undergo complete postbreeding moult, usually finished by September in great tits and by October in blue tits (Demongin 2016).

**Study site, monitoring and sampling**

Bielany Forest (ca. 150 ha) is situated in the northern part of the city of Warsaw and its northern border adjoins the left bank of the Vistula River. The area is a remnant of the Mazovian primeval forest and due to its unique natural value, it is protected under the Natura 2000 network (PLH140041, Special Area of Conservation, 129.84 ha) and is under reserve protection (as the Nature Reserve Bielany Forest, 130.35 ha). Bielany Forest is inhabited by ca. 1000 invertebrate species (including rare beetles - *Cerambyx cerdo* and *Osmoderma eremita*), 20 species of mammals and 40 breeding species of birds (Pawłat-Zawrzykraj et al. 2021). Because of its connection with Kampinos National Park (via Młociny Park and Forest and Vistula River valley) it plays a role as important ecological corridor for bigger mammals, e.g., wild boars, roe and red deers, foxes and hares.

Our study was conducted over two consecutive breeding seasons (2018-2019) within two plots: natural-cavity and nestbox, separated by a local road and within a minimum of 200 m of each other (Fig S1). The natural-cavity plot (52°17'32.2"N 20°57'34.0"E) consisted of 50 ha of monitored area (with 30 ha of core area with most intense nest searches) and was characterised by a very high availability of tree hollows, as only ca. 21% of cavities occupied in 2019 were used in previous breeding seasons. The nestbox plot (52°17'52.1"N 20°57'09.7"E) had an area of 15 ha and consisted of 65 nestboxes (woodcrete Schwegler 1b, Fig S5) hung every 50 meters, at the medium height of 2.91 m. The Schwegler nestboxes have an entrance hole of 32 mm of diameter, 12 cm of internal diameter and were 24 cm high. For details on basic microclimate variables in these nestboxes in comparison to plywood nestboxes and natural cavities, please refer to Sudyka et al. 2022a, and for more information on food availability, noise and air pollution within the forest see Sudyka et al. 2022b.

Nest searches in the natural cavity plot and nestbox monitoring started at the beginning of April and were continued throughout the season (to find potential new nests of pairs that experienced brood failure). The identified nests were monitored to record laying date, clutch size, incubation start, hatching date, number of fledglings and fledging date. Nestlings underwent breast feathers sampling and biometric measurements at 14^th^ day after hatching (hatching day = day 0). Adults were caught with mist nets or traps installed inside nest-boxes, ca. 14^th^ day after chicks hatching. Sex and age of adults were assigned according to the presence of the brood patch and moult limit between the primary and greater coverts, respectively (Svensson 1994). We measured body mass, tarsus length and wing length as well as took feather samples: crown and breast feathers from blue tits and breast feathers from great tits. Feather samples were stuck to and preserved on a black paper covered with a double adhesive, transparent tape.

**Statistical analysis – adults colouration**

To infer potential differences in colour traits of adult individuals breeding in natural cavities and nestboxes, we ran separate general linear mixed models, treating brightness and carotenoid chroma of breast feathers and brightness and UV chroma for blue tit crown as a response variable (Table S4). The models included categorical variables of cavity type (natural cavity or nestbox), sex, age (individuals in their second year or older) and year (2018 or 2019) and random term of nest ID. In the initial models, we tested for three-way interactions between cavity type, sex and age, two-way interactions between cavity type and other categorical factors, as well as interactions between age and sex. Whenever non-significant (P < 0.05), interactions were sequentially removed from the models. Multicollinearity of predictors in final models was controlled by checking the variance inflation factor (VIF), and it was lower than 2 in all cases. Statistical analysis were performed in R [version 4.1.1, (R Core Team, 2021)].

**Tables**

**Table S1.** Sample size of blue tit and great tit nestlings, with number of nests given in parentheses.

|  | **Year** | | **Blue tit** | **Great tit** |
| --- | --- | --- | --- | --- |
| **Natural cavities** | | 2018 | 73 (9) | 100 (15) |
|  | | 2019 | 123 (17) | 87 (14) |
| **Nest boxes** | | 2018 | 175 (20) | 120 (14) |
|  | | 2019 | 96 (12) | 59 (10) |
| **Total** | |  | 467 (58) | 366 (53) |

**Table S2.** Results of full general linear mixed models examining the influence of cavity type, parents’ plumage characteristics and age, nestlings’ sex and mass, clutch size, laying date and year on the plumage traits of blue tit and great tit nestlings. To account for the impact of common rearing environment and the cases when nestlings were weighted on other than 14th day, we fitted a unique nest ID (nest ID with a year code, hereafter: nest ID) and Time shift D14 as a random terms. Interactions between cavity type and all other main factors were sequentially removed if not significant. We also tested for non-linearity of lay date effect by introducing lay date^2, but as it was never significant (all P < 0.1), we do not report it here. All colour variables and clutch size were scaled to zero mean and unit SD. The table shows estimates, standard errors, degrees of freedom, t-values, and p-values. P<0.05 are given in bold and P<0.1 in italics, *P<0.05, **P<0.01, ***P<0.001.

|  | **Estimate** | | **SE** | **df** | **t** | **p** |  |
| --- | --- | --- | --- | --- | --- | --- | --- |
| **BLUE TIT** | |  |  |  |  |  |  |
| *Brightness* | |  |  |  |  |  |  |
| Intercept | | -0.236 | 0.21 | 34.8 | -1.14 | 0.262 |  |
| Cavity type (Box) | | 0.272 | 0.17 | 37.01 | 1.6 | 0.118 |  |
| Female brightness | | -0.045 | 0.08 | 30.21 | -0.56 | 0.583 |  |
| Male brightness | | -0.035 | 0.08 | 37.28 | -0.43 | 0.668 |  |
| **Female age (after second year)** | | **0.421** | **0.16** | **35.13** | **2.71** | **0.01** | ***** |
| Male age (after second year) | | -0.124 | 0.16 | 35.64 | -0.77 | 0.449 |  |
| Clutch size | | -0.021 | 0.08 | 36.7 | -0.28 | 0.783 |  |
| Lay date | | 0.014 | 0.11 | 35.96 | 0.13 | 0.895 |  |
| Nestling' mass | | -0.078 | 0.07 | 172.47 | -1.18 | 0.239 |  |
| Nestling' sex (M) | | -0.139 | 0.11 | 306.99 | -1.23 | 0.221 |  |
| Year (2019) | | -0.034 | 0.19 | 35.47 | -0.17 | 0.863 |  |
| *Random effects:* | | Variance |  |  |  |  |  |
| Unique nest ID | | 0.093 |  |  |  |  |  |
| Residual | | 0.828 |  |  |  |  |  |
| *Carotenoid chroma* | |  |  |  |  |  |  |
| Intercept | | -0.36 | 0.80 | 207.11 | -0.45 | 0.654 |  |
| Cavity type (Box) | | 0.401 | 0.21 | 35.2 | 1.95 | *0.059* |  |
| Female brightness | | -0.067 | 0.11 | 31.99 | -0.62 | 0.542 |  |
| Male brightness | | 0.147 | 0.11 | 36.14 | 1.29 | 0.204 |  |
| Female age (after second year) | | -0.101 | 0.19 | 34.49 | -0.54 | 0.595 |  |
| Male age (after second year) | | -0.133 | 0.22 | 35.39 | -0.62 | 0.541 |  |
| **Clutch size** | | **-0.235** | **0.09** | **35.97** | **-2.77** | **0.009** | ****** |
| Lay date | | -0.187 | 0.12 | 35.27 | -1.52 | 0.137 |  |
| Nestling' mass | | 0.027 | 0.07 | 262.37 | 0.38 | 0.706 |  |
| **Nestling' sex (M)** | | **0.343** | **0.10** | **303.92** | **3.47** | **0.001** | ******* |
| Year (2019) | | -0.397 | 0.26 | 34.42 | -1.55 | 0.13 |  |
| *Random effects:* | | Variance |  |  |  |  |  |
| Unique nest ID | | 0.212 |  |  |  |  |  |
| Time_shift_D14 | | 0.034 |  |  |  |  |  |
| Residual | | 0.588 |  |  |  |  |  |
| **GREAT TIT** | |  |  |  |  |  |  |
| *Brightness* | |  |  |  |  |  |  |
| Intercept | | -1.135 | 0.71 | 106.07 | -1.6 | 0.112 |  |
| **Cavity type (Box)** | | **0.439** | **0.15** | **26.38** | **2.89** | **0.008** | ****** |
| Female brightness | | 0.102 | 0.07 | 25.75 | 1.42 | 0.167 |  |
| Male brightness | | -0.017 | 0.06 | 28.24 | -0.27 | 0.791 |  |
| Female age (after second year) | | 0.045 | 0.14 | 24.46 | 0.31 | 0.757 |  |
| Male age (after second year) | | 0.021 | 0.15 | 25.38 | 0.14 | 0.891 |  |
| **Lay date** | | **-0.192** | **0.07** | **28.6** | **-2.84** | **0.008** | ****** |
| Nestling' mass | | 0.051 | 0.05 | 121.5 | 1.12 | 0.266 |  |
| Nestling' sex (M) | | 0.181 | 0.12 | 220.49 | 1.56 | 0.12 |  |
| Year (2019) | | -0.119 | 0.14 | 26.69 | -0.84 | 0.409 |  |
| *Random effects:* | | Variance |  |  |  |  |  |
| Unique nest ID | | 0.031 |  |  |  |  |  |
| Residual | | 0.692 |  |  |  |  |  |
| *Carotenoid chroma* | |  |  |  |  |  |  |
| Intercept | | 0.258 | 0.87 | 185.68 | 0.3 | 0.768 |  |
| Cavity type (Box) | | 0.498 | 0.36 | 24.81 | 1.4 | 0.173 |  |
| Female brightness | | 0.085 | 0.12 | 21.15 | 0.72 | 0.479 |  |
| Male brightness | | -0.168 | 0.09 | 27.17 | -1.78 | *0.086* |  |
| Female age (after second year) | | 0.229 | 0.24 | 20.56 | 0.94 | 0.356 |  |
| Male age (after second year) | | 0.431 | 0.33 | 22.4 | 1.32 | 0.200 |  |
| **Clutch size** | | **0.361** | **0.12** | **23.88** | **3.09** | **0.005** | ****** |
| Lay date | | 0.18 | 0.11 | 20.23 | 1.69 | 0.106 |  |
| Nestling mass | | -0.049 | 0.06 | 202.75 | -0.89 | 0.377 |  |
| Nestling sex (M) | | 0.065 | 0.11 | 212.57 | 0.58 | 0.565 |  |
| Year (2019) | | 0.408 | 0.23 | 20.46 | 1.75 | *0.095* |  |
| **Cavity type: Male age** | | **-1.104** | **0.48** | **19.72** | **-2.31** | **0.032** | ***** |
| Random effects: | | Variance |  |  |  |  |  |
| Unique nest ID | | 0.231 |  |  |  |  |  |
| Time shift D14 | | 0.089 |  |  |  |  |  |
| Residual | | 0.617 |  |  |  |  |  |

**Table S3.** Results of full general linear mixed models performed on original data set, without removed outliers. The models examined the influence of cavity type, parents’ plumage characteristics and age, nestlings’ sex and mass, clutch size, laying date and year on the plumage traits of blue tit and great tit nestlings. Interactions between cavity type and all other main factors were sequentially removed if not significant. All colour variables and clutch size were scaled to zero mean and unit SD. The table shows estimates, standard errors, degrees of freedom, t-values, and p-values. P<0.15 are given in italics, *P<0.05, **P<0.01, ***P<0.001.

|  | **Estimate** | **SE** | **df** | **t** | **p** |  |
| --- | --- | --- | --- | --- | --- | --- |
| **BLUE TIT** |  |  |  |  |  |  |
| *Brightness* |  |  |  |  |  |  |
| Intercept | 0.835 | 0.797 | 176.851 | 1.047 | 0.296 |  |
| Cavity type (Box) | 0.218 | 0.170 | 38.954 | 1.284 | 0.207 |  |
| Female brightness | -0.005 | 0.085 | 33.666 | -0.057 | 0.955 |  |
| Male brightness | -0.067 | 0.086 | 38.145 | -0.782 | 0.439 |  |
| **Female age (older)** | **0.383** | **0.161** | **37.816** | **2.380** | **0.022** | ***** |
| Male age (older) | -0.176 | 0.171 | 38.719 | -1.034 | 0.308 |  |
| Clutch size | -0.038 | 0.090 | 39.209 | -0.428 | 0.671 |  |
| **Lay date** | **-0.210** | **0.097** | **49.545** | **-2.151** | **0.036** | ***** |
| Nestling' mass | -0.085 | 0.072 | 190.042 | -1.184 | 0.238 |  |
| Nestling' sex (M) | -0.098 | 0.111 | 317.879 | -0.882 | 0.378 |  |
| Year (2019) | -0.186 | 0.182 | 38.375 | -1.024 | 0.312 |  |
| *Carotenoid chroma* |  |  |  |  |  |  |
| Intercept | -0.811 | 0.763 | 223.889 | -1.062 | 0.289 |  |
| **Cavity type (Box)** | **0.416** | **0.191** | **38.705** | **2.180** | **0.035** | ***** |
| Female carotenoid chroma | -0.068 | 0.107 | 33.875 | -0.638 | 0.528 |  |
| Male carotenoid chroma | 0.149 | 0.111 | 38.808 | 1.343 | 0.187 |  |
| Female age (older) | -0.083 | 0.183 | 36.576 | -0.452 | 0.654 |  |
| Male age (older) | -0.113 | 0.211 | 38.868 | -0.537 | 0.594 |  |
| **Clutch size** | **-0.259** | **0.092** | **38.454** | **-2.820** | **0.008** | ****** |
| **Lay date** | **-0.254** | **0.107** | **43.223** | **-2.371** | **0.022** | ***** |
| Nestling' mass | 0.060 | 0.069 | 270.150 | 0.867 | 0.387 |  |
| **Nestling' sex (M)** | **0.327** | **0.098** | **314.483** | **3.339** | **0.001** | ******* |
| Year (2019) | -0.373 | 0.234 | 36.211 | -1.596 | 0.119 |  |
| **GREAT TIT** |  |  |  |  |  |  |
| *Brightness* |  |  |  |  |  |  |
| Intercept | -0.592 | 0.724 | 127.864 | -0.817 | 0.415 |  |
| **Cavity type (Box)** | **0.473** | **0.143** | **23.179** | **3.305** | **0.003** | ****** |
| Female brightness | 0.120 | 0.066 | 22.369 | 1.804 | *0.085* |  |
| **Male brightness** | **-0.255** | **0.082** | **45.349** | **-3.097** | **0.003** | ****** |
| Female age (older) | 0.112 | 0.137 | 21.126 | 0.812 | 0.426 |  |
| Male age (older) | -0.021 | 0.139 | 23.819 | -0.153 | 0.880 |  |
| Clutch size | 0.019 | 0.072 | 31.221 | 0.262 | 0.795 |  |
| **Lay date** | **-0.186** | **0.066** | **25.668** | **-2.805** | **0.009** | ****** |
| Nestling' mass | 0.015 | 0.046 | 145.740 | 0.331 | 0.741 |  |
| Nestling' sex (M) | 0.188 | 0.108 | 221.673 | 1.730 | *0.085* |  |
| Year (2019) | -0.118 | 0.142 | 23.358 | -0.832 | 0.414 |  |
| **Cavity type: Male Brightness** | **0.360** | **0.124** | **28.359** | **2.909** | **0.007** | ****** |
| *Carotenoid chroma* |  |  |  |  |  |  |
| Intercept | 0.483 | 0.875 | 188.244 | 0.552 | 0.581 |  |
| Cavity type (Box) | 0.370 | 0.368 | 25.240 | 1.007 | 0.324 |  |
| Female carotenoid chroma | -0.031 | 0.128 | 19.811 | -0.238 | 0.814 |  |
| Male carotenoid chroma | 0.023 | 0.128 | 22.200 | 0.183 | 0.856 |  |
| **Female age (older)** | **0.314** | **0.125** | **21.157** | **2.513** | **0.020** | ***** |
| Male age (older) | 0.159 | 0.113 | 19.363 | 1.402 | 0.177 |  |
| Clutch size | 0.179 | 0.262 | 18.208 | 0.683 | 0.503 |  |
| Lay date | 0.554 | 0.349 | 23.309 | 1.586 | 0.126 |  |
| Nestling' mass | -0.062 | 0.056 | 205.426 | -1.097 | 0.274 |  |
| Nestling' sex (M) | 0.086 | 0.113 | 211.732 | 0.756 | 0.451 |  |
| Year (2019) | 0.403 | 0.277 | 20.567 | 1.458 | 0.160 |  |
| **Cavity type: Male Age** | **-1.114** | **0.506** | **20.086** | **-2.202** | **0.040** | ***** |

**Table S4.** Summary statistics of linear mixed effect models examining the relationship between adult blue tit and great tit plumage characteristics and cavity type (where the natural cavity is set as reference value to the nestbox effect), year, age (first year breeding bird or older), and sex. Interactions between cavity type and P<0.1 are given in italics, *P<0.05, **P<0.01, ***P<0.001.

|  | **Estimate** | **SE** | **df** | **t** | **p** |  |
| --- | --- | --- | --- | --- | --- | --- |
| **BLUE TIT** |  |  |  |  |  |  |
| *Crown brightness* |  |  |  |  |  |  |
| Intercept | -0.271 | 0.19 | 92.11 | -1.395 | 0.166 |  |
| Cavity type (nestbox) | 0.236 | 0.17 | 62.96 | 1.404 | 0.165 |  |
| **Year (2019)** | **-0.586** | **0.17** | **63.56** | **-3.473** | **0.001** | ******* |
| Age (older) | -0.035 | 0.16 | 108.77 | -0.216 | 0.829 |  |
| **Sex (M)** | **0.903** | **0.15** | **64.09** | **6.043** | **<0.001** | ******* |
| *Random effects:* | Variance |  |  |  |  |  |
| Nest ID | 0.084 |  |  |  |  |  |
| Residual | 0.603 |  |  |  |  |  |
| *Crown UV chroma* |  |  |  |  |  |  |
| Intercept | -0.959 | 0.17 | 95.43 | -5.697 | 0.000 | *** |
| Cavity type (nestbox) | -0.241 | 0.13 | 59.13 | -1.794 | *0.078* |  |
| **Year (2019)** | **1.435** | **0.13** | **60.17** | **11.427** | **<0.001** | ******* |
| **Age (older)** | **0.386** | **0.17** | **80.25** | **2.237** | **0.028** | ***** |
| **Sex (M)** | **0.743** | **0.19** | **108.70** | **3.923** | **<0.001** | ******* |
| **Cavity type (nestbox): Age (older)** | **-0.620** | **0.26** | **107.47** | **-2.408** | **0.018** | ***** |
| *Random effects:* | Variance |  |  |  |  |  |
| Nest ID | 0.025 |  |  |  |  |  |
| Residual | 0.430 |  |  |  |  |  |
| *Breast brightness* |  |  |  |  |  |  |
| Intercept | -0.060 | 0.23 | 91.31 | -0.261 | 0.795 |  |
| Cavity type (nestbox) | -0.294 | 0.21 | 62.71 | -1.421 | 0.160 |  |
| Year (2019) | -0.133 | 0.21 | 63.31 | -0.640 | 0.524 |  |
| Age (older) | 0.086 | 0.19 | 109.96 | 0.455 | 0.650 |  |
| **Sex (M)** | **0.470** | **0.16** | **60.26** | **2.882** | **0.005** | ****** |
| *Random effects:* | Variance |  |  |  |  |  |
| Nest ID | 0.244 |  |  |  |  |  |
| Residual | 0.712 |  |  |  |  |  |
| *Breast carotenoid chroma** |  |  |  |  |  |  |
| Intercept | 0.376 | 0.22 | 89.95 | 1.686 | 0.095 |  |
| Cavity type (nestbox) | -0.243 | 0.20 | 61.36 | -1.197 | 0.236 |  |
| **Year (2019)** | **-0.653** | **0.20** | **61.95** | **-3.204** | **0.002** | ****** |
| Age (older) | -0.143 | 0.18 | 109.89 | -0.791 | 0.431 |  |
| Sex (M) | 0.268 | 0.15 | 58.08 | 1.750 | 0.085 |  |
| *Random effects:* | Variance |  |  |  |  |  |
| Nest ID | 0.282 |  |  |  |  |  |
| Residual | 0.614 |  |  |  |  |  |
| **GREAT TIT** |  |  |  |  |  |  |
| *Breast brightness* |  |  |  |  |  |  |
| Intercept | -0.452 | 0.19 | 82.83 | -2.442 | 0.017 | * |
| Cavity type (nestbox) | 0.214 | 0.21 | 58.86 | 1.038 | 0.304 |  |
| Year (2019) | -0.147 | 0.20 | 90.83 | -0.732 | 0.466 |  |
| **Age (older)** | **0.706** | **0.19** | **55.74** | **3.810** | **<0.001** | ******* |
| Sex (M) | 0.242 | 0.20 | 56.22 | 1.229 | 0.224 |  |
| *Random effects:* | Variance |  |  |  |  |  |
| Nest ID | 0.042 |  |  |  |  |  |
| Residual | 0.852 |  |  |  |  |  |
| *Breast carotenoid chroma* |  |  |  |  |  |  |
| Intercept | 0.223 | 0.20 | 70.02 | 1.138 | 0.259 |  |
| Cavity type (nestbox) | 0.055 | 0.29 | 56.93 | 0.191 | 0.850 |  |
| Year (2019) | -0.395 | 0.27 | 54.30 | -1.466 | 0.148 |  |
| Age (older) | 0.274 | 0.20 | 94.99 | 1.381 | 0.171 |  |
| Sex (M) | -0.092 | 0.16 | 50.33 | -0.586 | 0.560 |  |
| **Cavity type (nestbox): Year (2019)** | **-0.912** | **0.43** | **56.87** | **-2.124** | **0.038** | ***** |
| *Random effects:* | Variance |  |  |  |  |  |
| Nest ID | 0.246 |  |  |  |  |  |
| Residual | 0.591 |  |  |  |  |  |

**Figures**

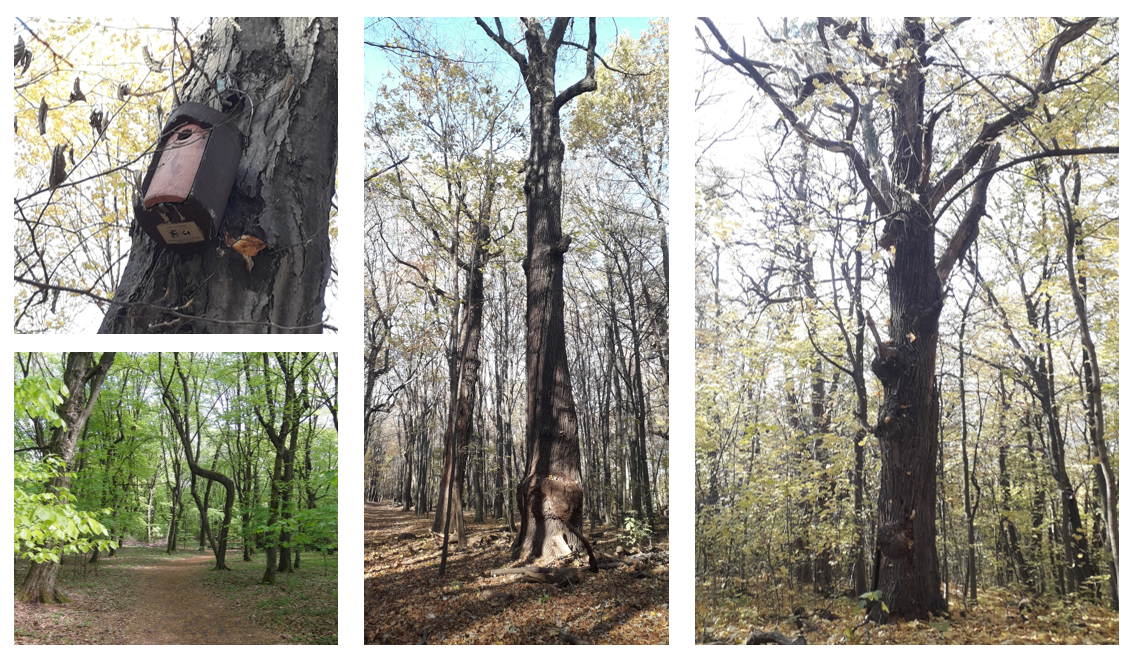

**Figure S1.** Overview of the Bielany Forest, with Schwegler’s nestbox in the upper left panel (photos by Piotr Janas and Joanna Sudyka).

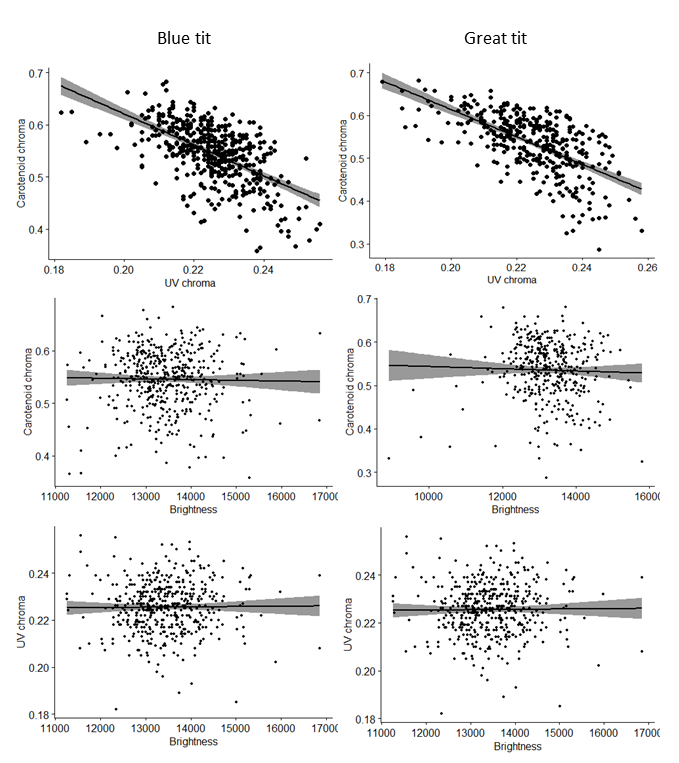

**Figure S2.** Relationships between blue tit (left panel) and great tit (right panel) nestlings breast brightness, carotenoid chroma and UV chroma Black line represents regression line and grey area represents confidence intervals.

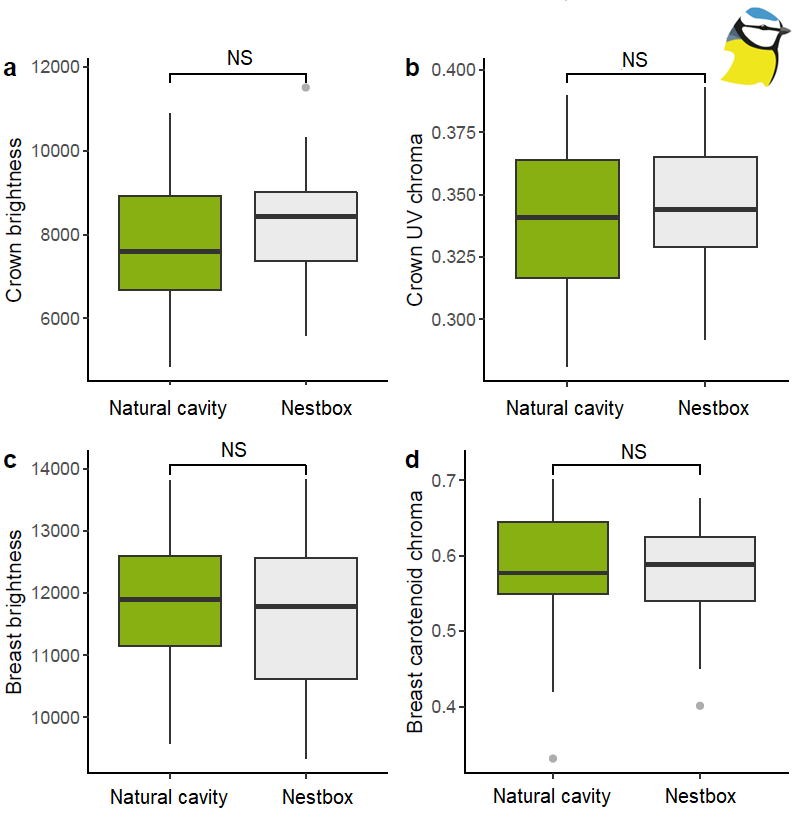

**Figure S3.** Boxplots of adult blue tits crown brightness (a), crown UV chroma (b), breast brightness (c) and breast carotenoid chroma (d). Sample size: 119 blue tits (61 females and 58 males) from 58 natural cavities and 61 nestboxes. Black horizontal bars denote raw data median, whiskers indicate minimum and maximum values, dark green and light grey colours denote, respectively, adults breeding in natural cavities and nest-boxes. NS indicate lack of significant difference.

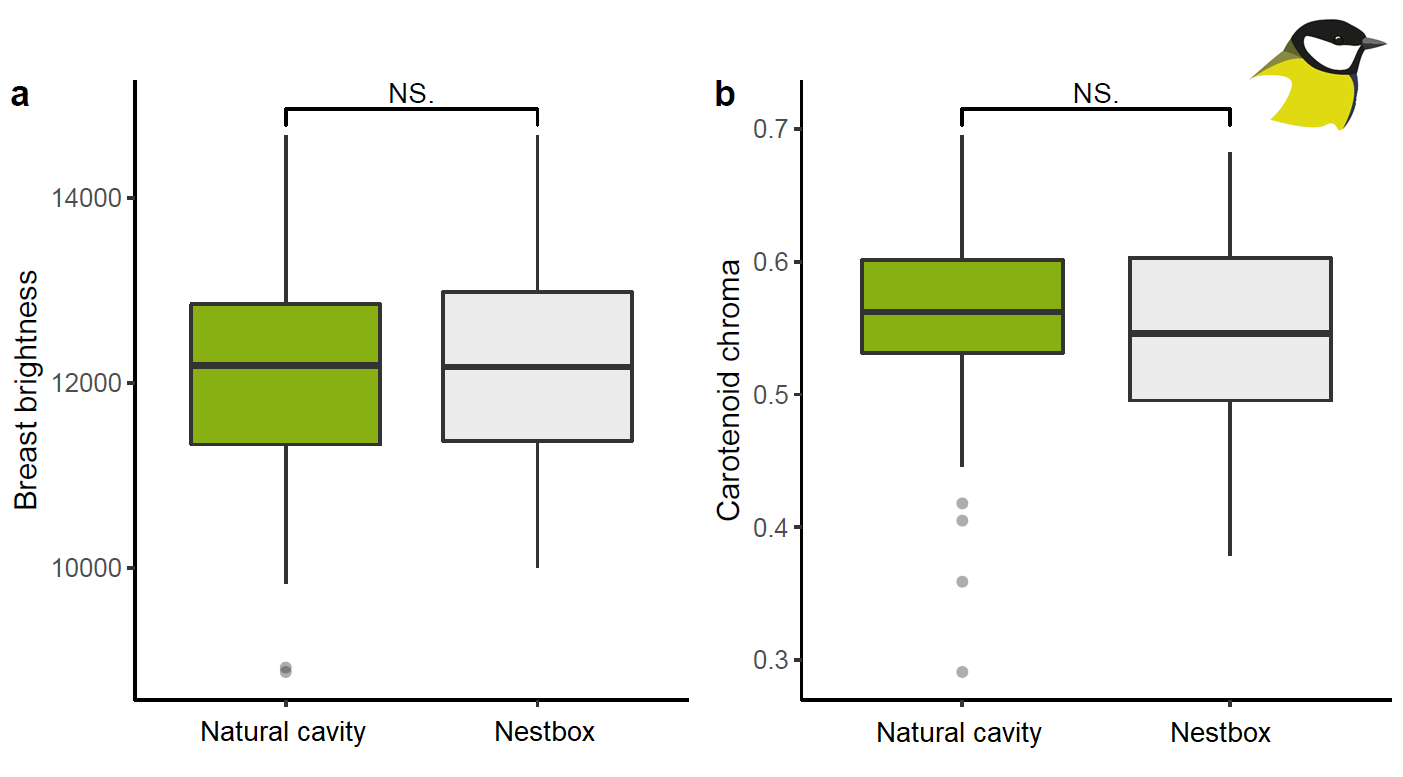

**Figure S4**. Boxplots of adult great tits breast brightness (a) and carotenoid chroma (b). Black horizontal bars denote raw data median, whiskers indicate minimum and maximum values, dark green and light grey colours denote, respectively, adults breeding in natural cavities and nest-boxes. Sample size: 115 great tits (61 females and 54 males) from 66 natural cavities and 49 nestboxes. NS indicate lack of significant difference.

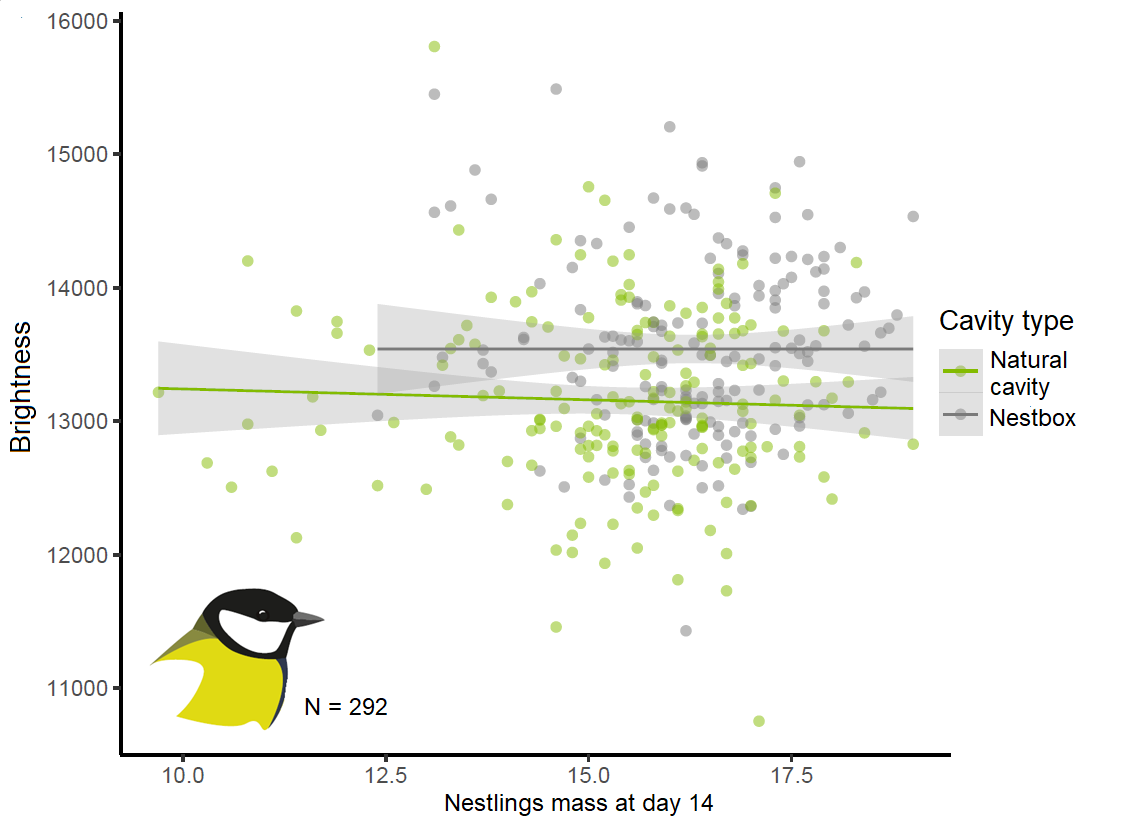

**Figure S5.** Scatterplot of great tit nestlings breast brightness as predicted by nestlings’ mass at day 14. Green points and line represent nestlings from natural cavities and grey points and line denote nestlings raised in nestboxes. Raw data with outliers removed.
